# Hepatocyte Nuclear Factor 4 alpha (HNF4α) Activation is Essential for Termination of Liver Regeneration

**DOI:** 10.1101/304808

**Authors:** Ian Huck, Sumedha Gunewardena, Regina Espanol-Suner, Holger Willenbring, Udayan Apte

**Affiliations:** Department of Pharmacology, Toxicology and Therapeutics, and University of Kansas Medical Center, Kansas City, KS; Department of Biostatistics University of Kansas Medical Center, Kansas City, KS; Eli and Edythe Broad Center of Regeneration Medicine and Stem Cell Research, University of California San Francisco, San Francisco, CA; Liver Center and University of California San Francisco, San Francisco, CA; Department of Surgery, Division of Transplantation, University of California San Francisco, San Francisco, CA

## Abstract

Hepatocyte Nuclear Factor 4 alpha (HNF4α) is critical for hepatic differentiation. Recent studies have highlighted its role in inhibition of hepatocyte proliferation and tumor suppression. However, the role of HNF4α in liver regeneration is not known. We hypothesized that hepatocytes modulate HNF4α activity when navigating between differentiated and proliferative states during liver regeneration. Western blot analysis revealed a rapid decline in nuclear and cytoplasmic HNF4α protein levels accompanied with decreased target gene expression within 1 hour after 2/3 partial hepatectomy (post-PH) in C57BL/6J mice. HNF4α protein expression did not recover to the pre-PH levels until day 3. Hepatocyte-specific deletion of HNF4α (HNF4α-KO) in mice resulted in 100% mortality post-PH despite increased proliferative marker expression throughout regeneration. Sustained loss of HNF4α target gene expression throughout regeneration indicated HNF4α-KO mice were unable to compensate for loss of HNF4α transcriptional activity. Deletion of HNF4α resulted in sustained proliferation accompanied by c-myc and cyclin D1 over expression and a complete deficiency of hepatocyte function after PH. Interestingly, overexpression of degradation-resistant HNF4α in hepatocytes did not prevent initiation of regeneration after PH. Finally, AAV8-mediated reexpression of HNF4α in hepatocytes of HNF4α-KO mice post-PH restored HNF4α protein levels, induced target gene expression and improved survival of HNF4α-KO mice post-PH. In conclusion, these data indicate that HNF4α reexpression following initial decrease is critical for hepatocytes to exit from cell cycle and resume function during the termination phase of liver regeneration. These results reveal the role of HNF4α in liver regeneration and have implications for therapy of liver failure.

## Introduction

HNF4α is known as the master regulator of hepatocyte differentiation due its role in establishing hepatocyte-specific gene expression during embryonic development (1, 2), stabilizing the hepatic transcription factor network (3) and maintaining hepatocyte function (4). HNF4α regulates genes involved in xenobiotic metabolism, carbohydrate metabolism, fatty acid metabolism, bile acid synthesis, blood coagulation, and ureagenesis (5). Expression of HNF4α induces hepatocyte-like characteristics in induced pluripotent stem cells (6) and forced expression of HNF4α can be used to differentiate and decrease cancer progression in hepatocellular carcinomas (HCC) and hepatoma cell lines (7, 8). Furthermore, decreased HNF4α expression leads to loss of hepatic function and causes decompensation in cirrhotic rats (9).

Recent studies from our laboratory and others have revealed anti-proliferative properties of HNF4α (10, 11). HNF4α is anti-proliferative in kidney cells (12) and pancreatic β-cells (13). Furthermore, hepatocyte-specific deletion of HNF4α (HNF4α-KO) in mice results in spontaneous hepatocyte proliferation (10, 11) and promotes formation of diethylnitrosamine-induced HCCs (14).

During liver regeneration after 2/3 partial hepatectomy (PH), the most widely used model to study liver regeneration, multiple redundant mechanisms regulate initiation and termination of hepatocyte regeneration (15). Understanding the mechanisms that govern adult hepatocytes to navigate between quiescent and proliferative states could result in therapeutic targets for inducing hepatocyte proliferation during impaired regeneration or inhibiting excess proliferation during carcinogenesis. Despite its role in maintaining hepatocyte differentiation and quiescence, little is known about the role of HNF4α in hepatocyte regeneration or how decreased HNF4α, a condition commonly found in diseased human livers (9, 16–18), would impact regeneration. In this study, we investigated the role of HNF4α in regulation of liver regeneration after PH using wildtype (WT) and HNF4α-KO mice. Our studies revealed that HNF4α is indispensible for survival after PH and a critical component of termination of liver regeneration.

## Materials and Methods

### Animal Care and Surgeries

Animals were housed in facilities accredited by the Association for Assessment and Accreditation of Laboratory Animal Care at the University of Kansas Medical Center under a standard 14-hr dark/10-hr light cycle with access to chow and water *ad libitum.* All studies were approved by the Institutional Animal Care and Use Committee at the University of Kansas Medical Center. PH surgeries were performed and tissue samples obtained as previously described (19). Surgeries were performed between 9:00 am and 11:00 am. For liver regeneration in normal conditions, two-month-old male C57BL6/J mice were euthanized at 0, 1, 3, 6, 12, 24 and 48 hr, and 3, 5, 7, and 14 days after PH with 3 mice per time point. To study the effects of HNF4α deletion on liver regeneration, seven days before surgery HNF4α-floxed mice were injected intraperitoneally with AAV8-TBG-eGFP or AAV8-TBG-CRE resulting in WT and hepatocyte-specific HNF4α-KO animals, respectively. These vectors were purchased from Penn Vector Core (Philadelphia, PA) and injected as previously described(20). WT and HNF4α-KO mice were euthanized at 0, 1, 2, 5, 7 and 14 days after PH with 3-5 mice per group. Only two HNF4α-KO mice survived to be included at the 7 day time point. HNF4α reexpression was accomplished by tail vein injection of 2 × 10^11^ viral genomes/mouse of AAV8-CMV-HNF4α generated in the laboratory of Dr. Holger Willenbring. Male HNF4α-floxed mice were injected with AAV8-TBG-CRE to induce HNF4α deletion and 7 days later were injected with AAV8-CMV-HNF4α to restore HNF4α expression. Liver and serum samples were collected seven days after AAV8-CMV-HNF4α injection. Next, this system was used to test if HNF4α reexpression could rescue HNF4α-KO mice after partial hepatectomy. WT and HNF4α-KO mice underwent PH 7 days after injection with AAV8-TBG-eGFP or AAV8-TBG-CRE with four mice per treatment. Tail vein injection with reexpression vector or control saline occurred 2 days after surgery to allow for initiation of regeneration. Overexpression of HNF4α was accomplished using Tet-On-HNF4α transgenic mouse line developed at the KUMC Transgenic and Gene Targeting Institutional Facility. These mice contained a reverse tetracycline-controlled transcriptional activator which was constitutively expressed with a C/EBPβ promoter. In the presence of the tetracycline-derivative Doxycycline (Dox), this transcription factor binds a tetracycline response element (TRE) upstream of an shRNA-resistant HNF4α gene (kind gift from Dr. Stephen A. Duncan, Medical University of South Carolina). Tet-On-HNF4α were heterozygous for both transgenes. Doxycycline was administered for two weeks prior to surgery by replacing drinking water with a 1 mg/mL Dox in 5% Sucrose solution. Untreated mice were given 5% sucrose solution only. Two mice were used for the untreated 6 hour time point and three to five mice used for all other treatments and time points.

### Protein Isolation and Western blotting

Nuclear and cytoplasmic lysates and RIPA extracts were prepared and used for Western blot analysis as reported (21). Primary antibodies are provided in Supplemental Table 1. Densitometric analysis was performed using Image Studio Lite (LI-COR Biosciences).

### Real Time PCR

RNA isolation, conversion to cDNA and Real time PCR analysis was performed as previously described (22). RNA sequencing and bioinformatics analysis were performed as previously described(14). Genes of interest were identified based on RNA-Seq analysis comparing gene expression in livers from WT and hepatocyte-specific HNF4α-KO mice(14). Primer sequences are provided in Supplemental Table 2.

### Staining Procedures

Paraffin-embedded liver sections (4-μm thick) were used for immunohistochemical staining of proliferating cell nuclear antigen (PCNA) as previously described(19). PCNA positive hepatocyte nuclei were quantified by counting three 40x fields per slide for each liver sample.

### Serum Bilirubin

Total serum bilirubin was determined using the Total Bilirubin kit from Pointe Scientific (Cat #B7576) according to manufacturer’s protocol.

### Statistical Analysis

Results are expressed as mean ±standard error. One Way ANOVA and Student’s *t* test was applied to all analyses with *p* < 0.05 being considered significant.

## Results

### Decreased HNF4α Protein Expression and Transcriptional Activity During Initiation of Regeneration After Partial Hepatectomy

First, we investigated the expression and activity of HNF4α in liver regeneration after PH in C57BL/6J mice. We measured protein expression of HNF4α adult isoforms using Western blotting in freshly prepared nuclear (Fig. 1A, 1C) and cytoplasmic (Fig. 1B) protein extracts. Nuclear HNF4α levels started declining at 1 (hour) hr post-PH reaching the lowest level at 6 hr post-PH. Nuclear HNF4a protein expression started rising at 12 hr, increased higher than the 0 hr levels peaking at 3 days post PH and then declined again to reach the 0 hr levels by 7 days post-PH (Fig. 1A, 1C). Cytoplasmic HNF4α expression continually declined below 0 hr levels until 12h post-PH, then started to rise but did not return to 0 hr pre-PH levels till 14 days after PH. The expression of adult isoform of HNF4α mRNA did not change over the entire regeneration time course (Fig. 1D).

**Figure 1.**
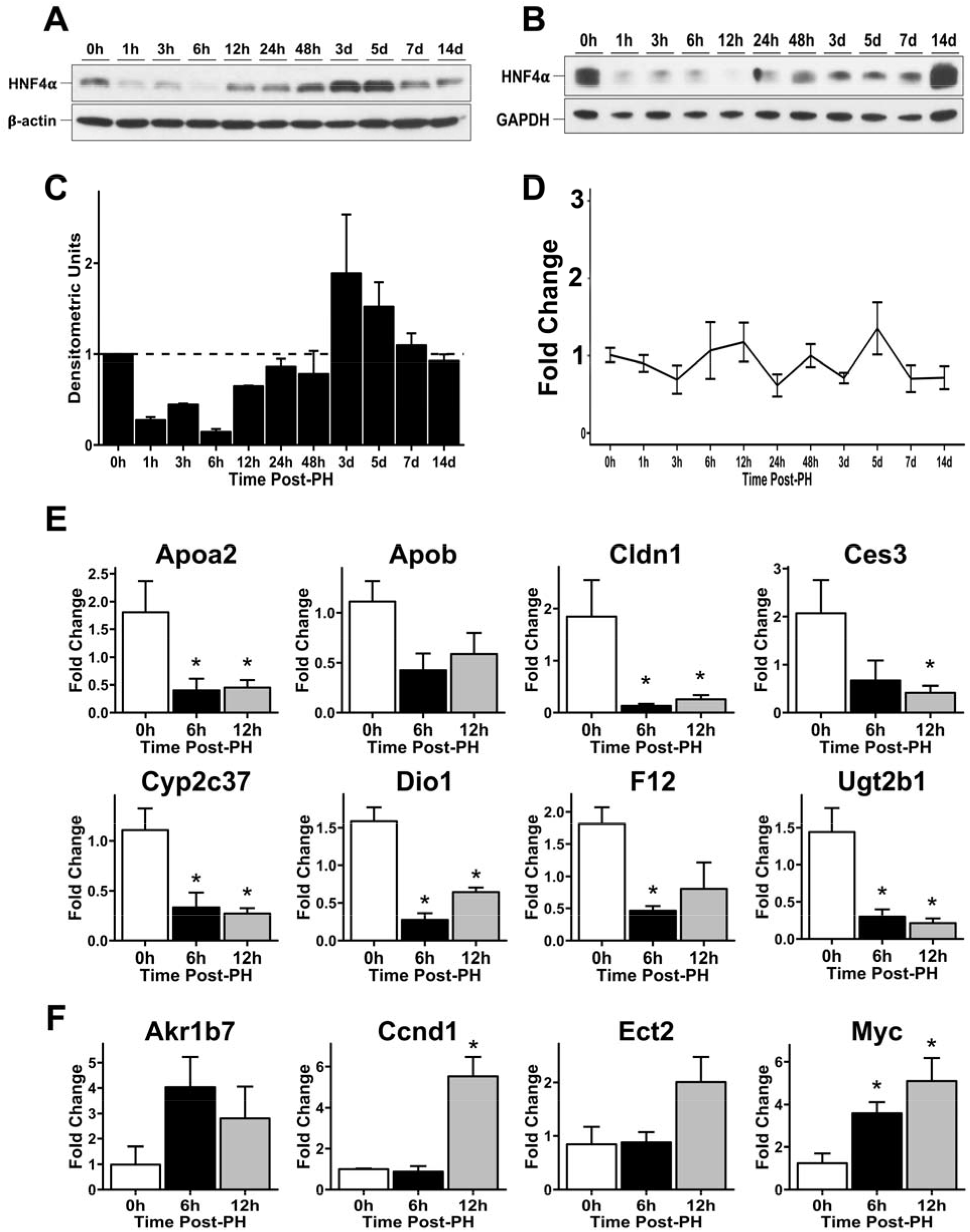
Decreased HNF4α protein expression and transcriptional activity during initiation of regeneration after partial hepatectomy. Western blot analysis of HNF4α adult isoform in (A) nuclear and (B) cytoplasmic lysates from mouse liver over a time course of 0 – 14 days after 2/3 PH. (C) Densitometric analysis of nuclear HNF4α blot. (D) qPCR analysis of HNF4α adult isoform mRNA over a time course of 0 – 14 days after 2/3 PH. Fold change calculated by comparison to 0 hr time point. qPCR analysis of positively regulated (E) and negatively regulated (F) HNF4α target genes at 0, 6 and 12 hr post-PH. Fold change calculated by comparison to 0 hr time point. *indicate significant difference at P ≤ 0.05.

We assessed HNF4α transcriptional activity by qPCR analysis of its target genes at the 0, 6 and 12 hr time points when nuclear HNF4α protein levels were at their lowest levels. We observed decreased expression of positive target genes (14) (Apoa2, ApoB, Ces3, Cldn1, Cyp2c37, Dio1, F12, Ugt2b1) at the 6 and 12-hr time points compared to 0 hr expression levels (Fig. 1E). We have previously identified novel negative targets of HNF4α, which are suppressed by HNF4α in the normal liver (14). qPCR analysis showed induction of HNF4α negative target genes (Akr1b7, Ccnd1, Ect2, Myc) at the 6 and 12 hr time points (Fig. 1F). Together, these data indicate a decrease in HNF4α activity at the onset of regeneration which is consistent with the changes we observed in HNF4α protein expression (Fig. 1A-C).

### HNF4α Overexpression Delays But Does Not Prevent Hepatocyte Proliferation During Liver Regeneration

We used a Tet-On-HNF4α transgenic mouse system (23) to overexpress HNF4α in hepatocytes to test if the anti-proliferative effects of HNF4α would prevent hepatocyte proliferation during the initiation of liver regeneration (Supp. Fig. 1). Western blot analysis confirmed increased HNF4α in nuclear lysates from livers of doxycycline (Dox)-treated Tet-On-HNF4α mice at 6 hours post-PH (Supp. Fig. 2A). This resulted in fewer PCNA-positive nuclei in Dox-treated mice 6 hours post-PH. However, no difference in PCNA staining was observed between untreated and Dox treated mice 48 hours post-PH (Supp. Fig. 2B). Interestingly, Dox treatment inhibited the occurrence of transient steatosis 48 hours post-PH (Supp. Fig 2C).

### Hepatocyte-Specific HNF4α-KO Mice Do Not Survive After PH

Next, we investigated the effect of hepatocyte-specific HNF4α deletion on liver regeneration after PH. WT and HNF4α-KO mice underwent PH and were euthanized at 1, 2, 5, 7 and 14 days after surgery. HNF4α protein remained undetectable in HNF4α-KO mice throughout the time course after PH (Fig 2A). We observed 100% mortality in the HNF4α-KO group by day 11 post-PH (Fig. 2B). Interestingly, the recovery of liver weight was similar between WT and HNF4α-KO groups until 5 days post PH (Fig. 2C). Serum bilirubin was significantly elevated in HNF4α-KO mice at days 1, 2 and 5 post-PH (Fig. 2D). While serum bilirubin in HNF4α-KO animals did decrease over time, this assay was only performed on surviving mice and most animals in this group had died at later time points. We investigated changes in HNF4α target gene expression in WT and HNF4α-KO mice at 7 days after PH, which revealed a significant decrease in expression of positive target genes including *Alas2, Apoa2, Apob, Cldn1, Cyp2c37, Dio1, F12 and Ugt2b1* (Fig 3A). Similarly, the expression of the negative targets *Akr1b7, CcnD1, Cdkn3, Defb1, Ect2, Egr1, Myc and Slc34a2* was increased at 7 days post-PH (Fig. 3B).

**Figure 2.**
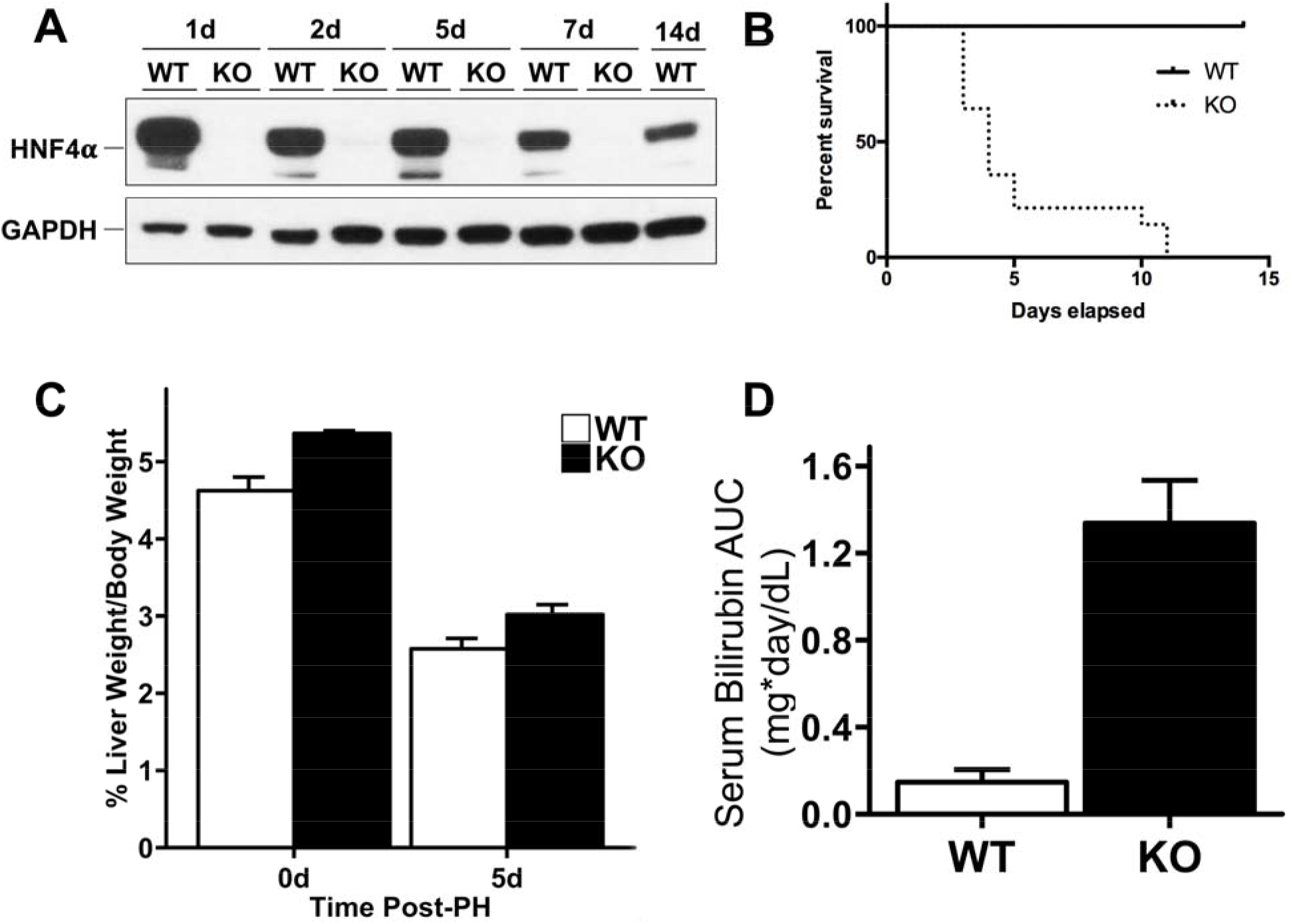
Complete mortality of HNF4α-KO mice following PH. (A) Western blot of HNF4α confirming efficient KO of HNF4α in pooled liver lysates at all time points post-PH. (B) Kaplan-Meier survival analysis of WT and HNF4α-KO groups after PH. (C) Liver weight to body weight ratios and (D) serum bilirubin levels in WT and HNF4α-KO mice after PH. *indicate significant difference at P ≤ 0.05 between WT and HNF4α-KO.

**Figure 3:**
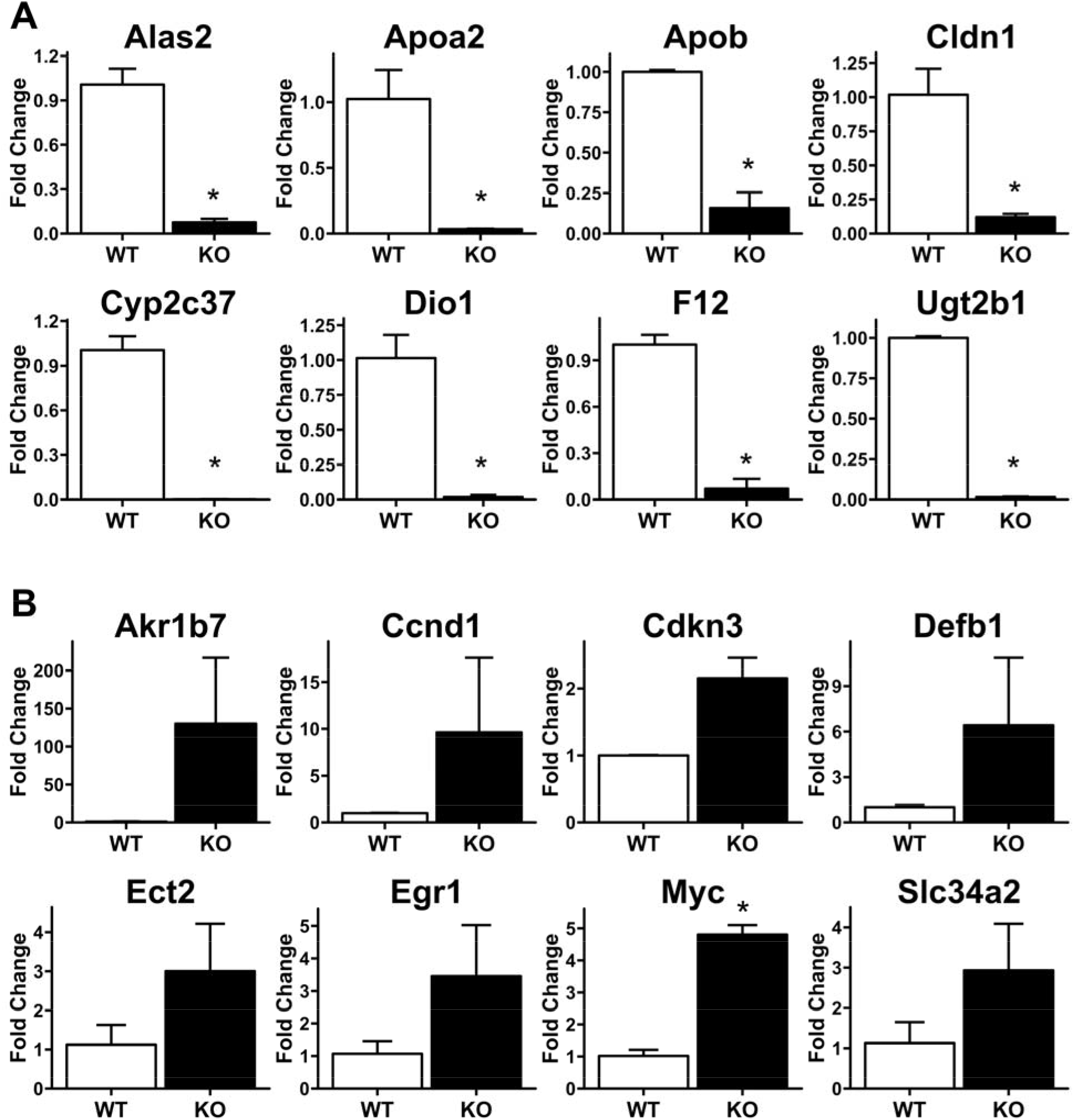
Sustained Loss of HNF4α Transcriptional Activity In HNF4α-KO Mice 7 Days Post-PH. qPCR analysis of mRNA isolated from frozen liver in WT and HNF4α-KO mice 7 days post-PH. (A) Decreased expression of positive targets of HNF4α and (B) increased expression of negative targets of HNF4α. *indicate significant difference at P ≤ 0.05 between WT and HNF4α-KO.

### Increased Hepatocyte Proliferation in HNF4α-KO Livers Throughout Regeneration

Western blot analysis indicated that protein expression of cyclin D1, the cyclin involved in driving G0 to G1 and G1 to S cell cycle transition, was significantly increased in HNF4α-KO livers at all time points. The expression of CDK4, the cyclin dependent kinase interacting with cyclin D1 was lower in HNF4α-KO liver at days 1 and 2 post PH but increased to levels comparable to WT mice by days 5 post PH. Cell proliferation assessed by Western blot and immunohistochemical analysis of PCNA revealed an elevated PCNA level in HNF4α-KO mice at all time points (Fig. 4A). Immunohistochemical analysis of PCNA-positive nuclei per 40x field was significantly higher in HNF4α-KO mice throughput the 7-day time course post PH (Fig. 4B-C).

**Figure 4.**
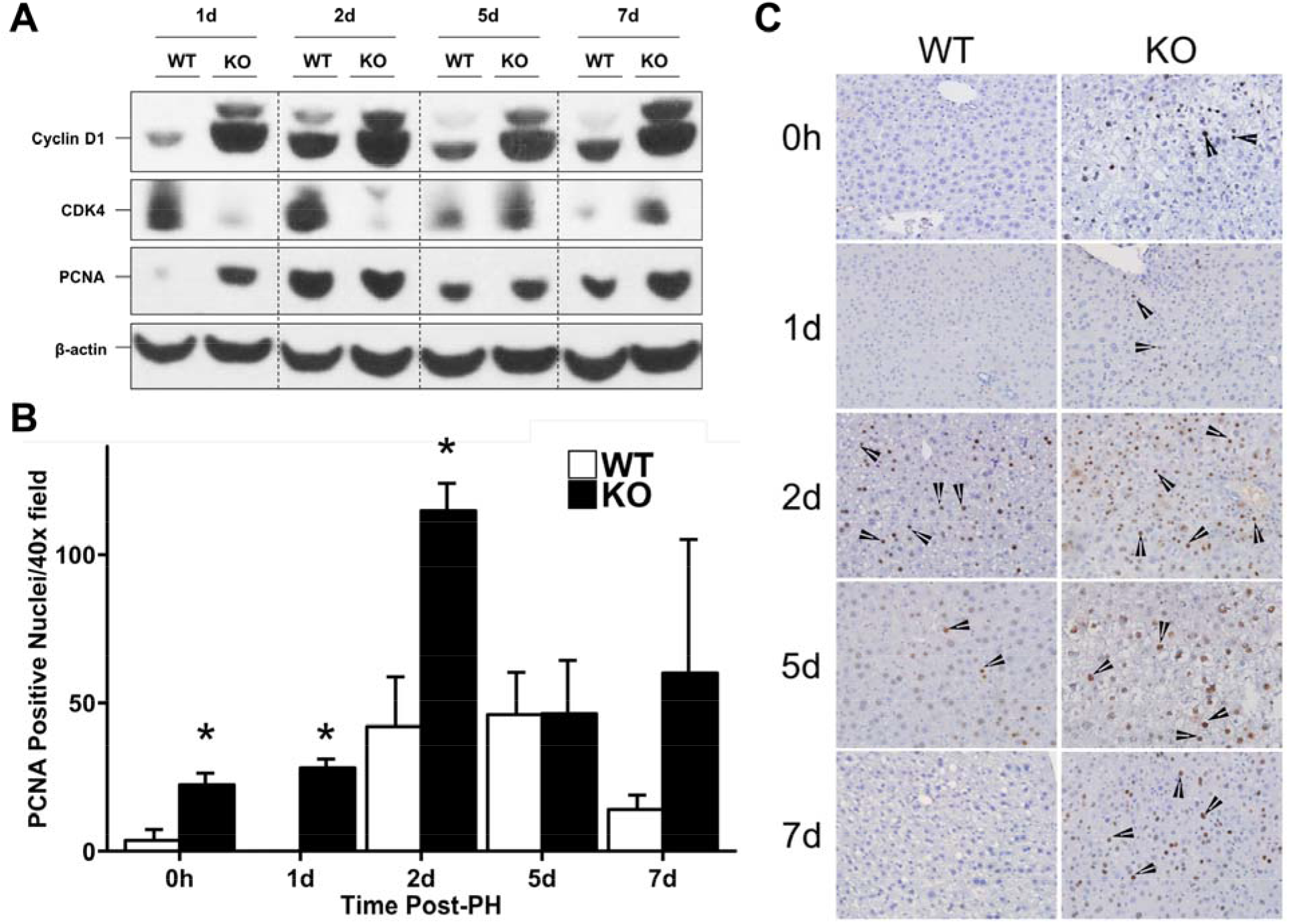
Increased hepatocyte proliferation in HNF4α-KO livers throughout regeneration. (A) Western blot analysis of Cyclin D1, CDK4, and PCNA over a time course of 0 to 7 days post-PH. (B) Quantification of immunohistochemical analysis for PCNA positive nuclei in liver sections of WT and HNF4α-KO mice throughout time course post-PH. (C) Representative photomicrographs (400x) of PCNA-stained liver sections from WT and HNF4α-KO mice throughout time course post-PH. *indicate significant difference at P ≤ 0.05 between WT and HNF4α-KO.

### Altered Proliferative Signaling in HNF4α-KO Mice During Regeneration

We investigated the mechanism of sustained hepatocyte proliferation throughout regeneration in HNF4α-KO mice by Western blot analysis of pathways commonly involved in hepatocyte proliferation. We focused on pathways downstream of primary hepatocyte mitogen signaling as well as those known to activate hepatocyte proliferation in the absence of HNF4α (14, 24). Most interestingly, we observed complete loss of total EGFR and total c-MET protein expression in HNF4α-KO animals at all time points throughout regeneration (Fig. 5A). Next, we examined Wnt/β-catenin pathway members (Fig. 5B). Total β-catenin levels were similar in WT and HNF4α-KO mice before PH and did not change throughout the time course. However, inactive (Thr41/Ser45-phosphorylated) β-catenin was elevated in the WT animals at days 1 and 2 post-PH. Phosphorylated β-catenin was greater in HNF4α-KO compared to WT mice 5 days post-PH. Densitometric analysis of inactive to total β-catenin ratio suggested activation of β-catenin was higher in HNF4α-KO mice at 1 and 2 days post-PH. Total GSK-3β levels did not change for both groups throughout the time course. Inactive (ser9-phospho) GSK-3β was higher in both groups at days 1 and 5 post-PH, but no difference was observed between groups. Next, we investigated activation of several MAPK members including AKT, p38 and ERK1/2 (Fig. 5C). No difference in total AKT and total p38 protein levels was observed between groups throughout regeneration. Phosphorylation of AKT was inhibited in the HNF4α-KO mice as compared to WT at days 0, 1 and 5 post-PH but was similar to WT at day 2 after PH. Phosphorylation (activation) of p38 was elevated in HNF4α-KO mice at days 1 and 5 post-PH. Finally, Total ERK1/2 expression was elevated in HNF4α-KO animals at all time points. Phosphorylation (activation) of ERK1 (upper band) was increased in HNF4α-KO animals compared to WT at all time points. Phosphorylation (activation) of ERK2 (lower band) was inhibited in HNF4α-KO group compared to WT at days 0 and 5 post-PH. ERK2 phosphorylation was elevated in HNF4α-KO group compared to WT at day 7 post-PH. Finally, we measured expression of known regulator of hepatocyte proliferation and negative target of HNF4α c-Myc (Fig. 5D). While c-Myc protein expression was similar between WT and HNF4α-KO animals at day 0, c-Myc expression was elevated in HNF4α-KO animals at all time points post-PH. Furthermore, c-Myc mRNA expression was elevated in HNF4α-KO mice at all time points.

**Figure 5.**
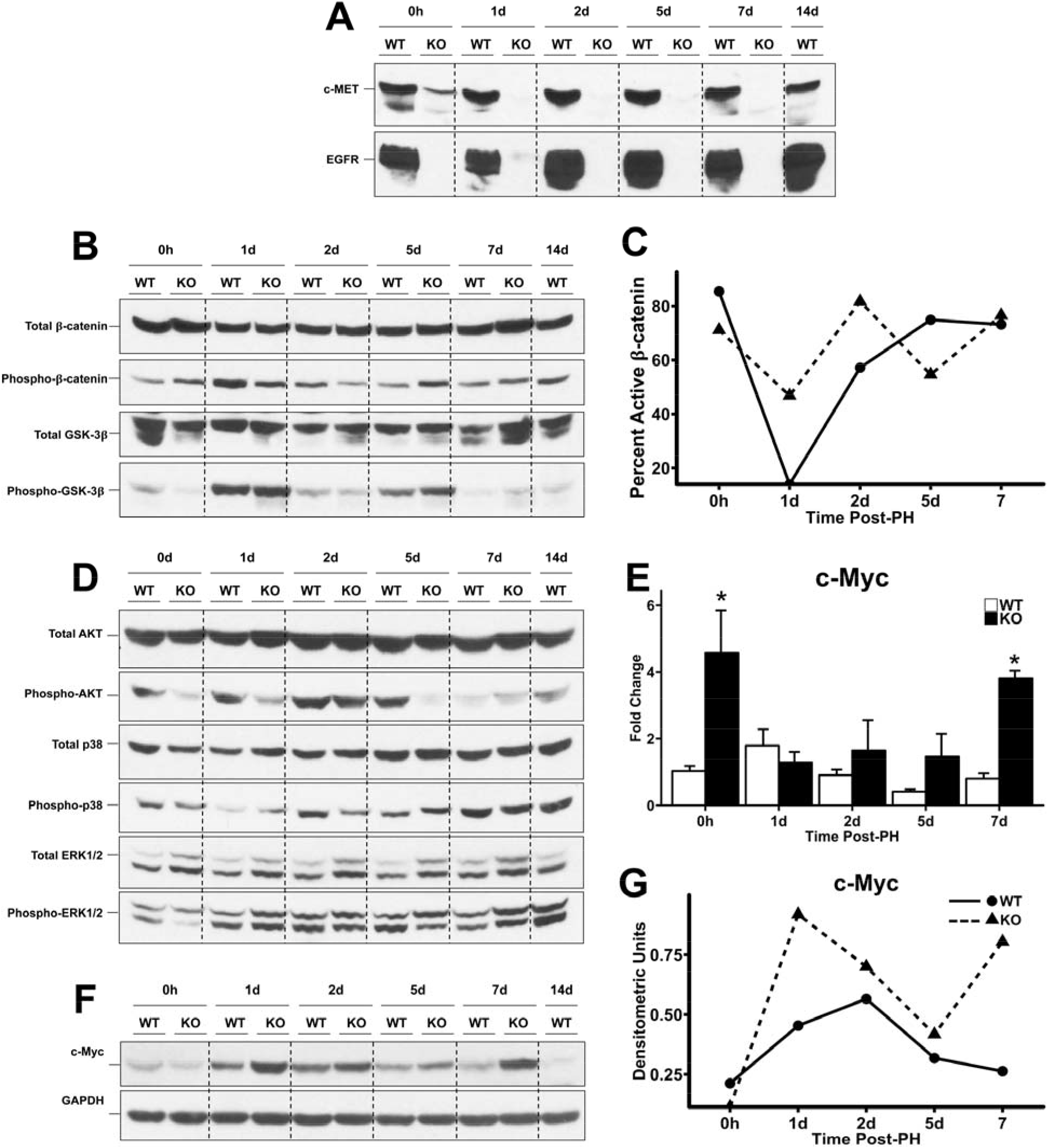
Altered proliferative signaling in HNF4α-KO mice during regeneration. Western blot analysis of (A) total EGFR and c-Met, (B) total and phosphorylated β-catenin and total and phosphorylatedGSK3β, (C) line graphs showing densitometric signal of inactive to total β-catenin, (D) Western blots for Total AKT, Phospho-AKT, Total p38, Phospho-p38, ERK1/2, Phospho-ERK1/2, (E) qPCR analysis of c-Myc mRNA from WT and HNF4α-KO livers over time course post-PH, (F) Western blot analysis of c-Myc, (G) and densitometric analysis of c-Myc. *indicate significant difference at P ≤ 0.05 between WT and HNF4α-KO.

### RNAseq revealed sustained increase in pro-proliferative and anti-differentiation signaling in HNF4α-KO Mice Post-PH

To gain insights in the comprehensive signaling changes in HNF4α-KO livers after PH, we performed RNA-Seq analysis at days 2 and 5 post-PH. These time points were selected to match the peak hepatocyte proliferation (day 2 post PH) and early termination phase of regeneration (day 5 post PH) in the liver regeneration process. Overall, a similar number of genes (~1,000) increased and decreased at day 2 and day 5 after PH in HNF4α-KO mice as compared to WT mice (Supp. Table 3). Ingenuity Pathway Analysis (IPA) predicted activation and inhibition of transcription factors based on gene expression changes between genotypes. At 2 days post-PH (Table 1), IPA predicted activation of effectors of proinflammatory signaling (EGR1, IRF3, IRF6, IRF7) and proproliferative transcription factors (JUN, RB1, P300, TCF3) in HNF4α-KO liver. Interestingly, HNF4α-KO also showed activation of TGF-β signaling as indicated by activation of SMAD2 and SMAD4 as well as inhibition of SMAD7 (25). The transcription factors predicted to be inhibited 2 days post-PH included HNF4α and its coactivators HNF1A and PPARGC1A (PGC1A). Furthermore, Estrogen Receptor α (Esrra), which is known to regulate gene expression in coordination with HNF4α and PGC1A (26), was also predicted to be inhibited. TAF4, a known HNF4α cofactor which is required for HNF4α transcriptional activity (27), was also inhibited. MED1, which is essential for PPARα activity and required for survival after PH (28), was inhibited in HNF4α-KO 2 days post-PH. ZBTB20, a known suppressor of hepatocyte proliferation and repressor of alpha-fetoprotein (29, 30), was predicted to be inhibited.

**Table 1:** Predicted Activity of Transcription Factors in HNF4α-KO Mice Compared to WT Mice 2 Days Post-PH.

| **Day 2 Post-PH** |  |  |  |
| --- | --- | --- | --- |
| **Activated Pathways** | **Activation z-score** | **Inhibited Pathways** | **Activation z-score** |
| JUN | 3.31 | HNF4A | -6.498 |
| EGR1 | 2.814 | HNF1A | -4.428 |
| RB1 | 2.497 | Esrra | -3.357 |
| IRF3 | 2.42 | PPARGC1A | -3.228 |
| FOXO4 | 2.377 | NFIX | -2.433 |
| EP300 | 2.305 | ZBTB20 | -2.294 |
| SPI1 | 2.248 | MED1 | -2.272 |
| THRAP3 | 2.219 | CLOCK | -2.219 |
| IRF6 | 2.215 | NFIC | -2.2 |
| SMAD4 | 2.201 | GATA2 | -2.059 |
| MTA1 | 2.194 | NKX2-3 | -2.04 |
| Gm21596/Hmgb1 | 2.169 | E2F1 | -2.027 |
| IRF7 | 2.163 | SMAD7 | -2.027 |
| SMAD2 | 2.068 | TAF4 | -2.008 |
| TCF3 | 2.022 |  |  |

At day 5 post-PH (Table 2), the analysis predicted activation of transcription factor proinflammatory damage associated molecular pattern (DAMP) protein HMGB1 in HNF4α-KO mice indicating sustained inflammatory signaling in HNF4α-KO mice throughout regeneration. The TGFβ effector SMAD2 was also activated at day 5 post-PH. Proliferative marker EP300 was activated at day 5 post-PH. The HNF4α negative target gene and proliferative marker CCND1 was activated in the HNF4α-KO group at day 5 post-PH indicated sustained proliferative signaling compared to WT mice. Activation of SNAI2 was predicted in the HNF4α-KO group at 5 days post-PH. SNAI2 is repressed by HNF4α and is known to promote EMT (31). PLAG1 is a fetal gene overexpressed in hepatoblastomas (32) and was predicted to be activated in the HNF4α-KO group at day 5. Activation of TRIM24, a transcription factor with oncogenic activity (33), was predicted at day 5 post PH in the HNF4α-KO mice. Transcription factors including HNF4A, HNF1A, PPARGC1A, and Esrra were inhibited at day 2 post-PH and continued to be inhibited at day 5 post-PH. Inhibition of SIRT2 was observed at 5 days post-PH in HNF4α-KO mice consistent with its known role in sharing many of the same target genes as HNF4α(34). PPARGC1B is a known coactivator of PPARGC1A and was inhibited. NCOA2 and EBF1, known tumor suppressors(35, 36), were inhibited in HNF4α-KO mice day 5 post-PH. NFIX, a transcription factor known to inhibit development of HCC(37), was inhibited. Finally, the master regulators of sterol and lipid metabolism, SREBF1 and SREBF2, were inhibited 5 days post-PH.

**Table 2:**
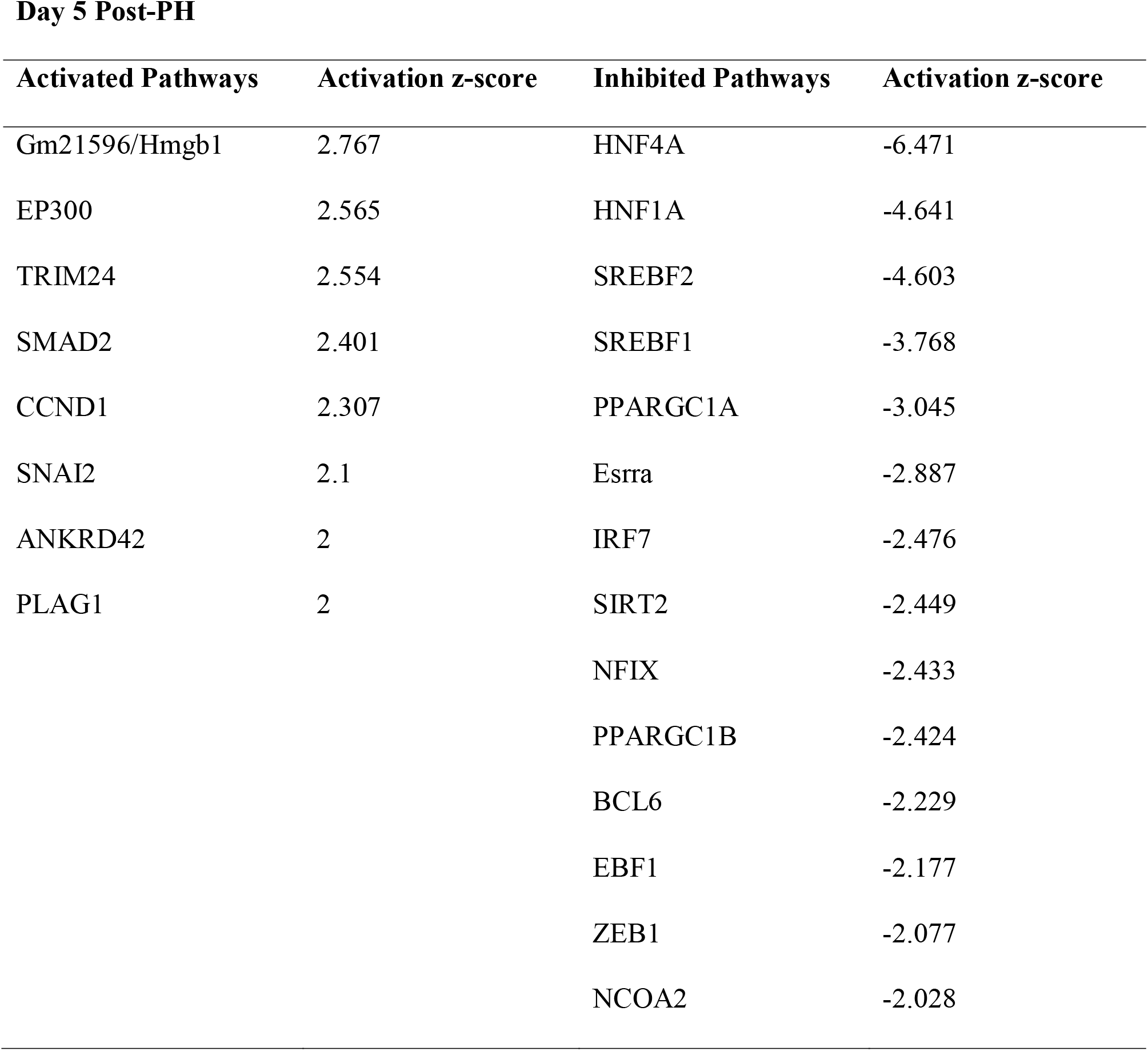
Predicted Activity of Transcription Factors in HNF4α-KO Mice Compared to WT Mice 5 Days Post-PH.

IPA analysis also predicted activity of diseases and organ function based on gene expression differences between WT and HNF4α-KO mice at 2 and 5 days post-PH. The comparison at day 2 post-PH (Supp. Table 4) predicted HNF4α-KO mice would exhibit activation of pathways involving inflammation, tumorigenesis and wound healing. Inhibited functions included numerous basic liver processes such as transport, metabolism and synthesis of cholesterol, lipids, bile acids and xenobiotics. This pattern continued at day 5 post-PH (Supp. Table 5). Activated functions continued to be related to cell proliferation, tumorigenesis and inflammation. Additionally, functions related to embryonic organ tissue development were activated in the HNF4α-KO group. Functions related to transport and metabolism of lipids, cholesterol and vitamins remained inhibited in HNF4α-KO mice 5 days post-PH.

### Reexpression of HNF4α Restores Hepatocyte Quiescence and Gene Expression and Extends Survival of HNF4α-KO Animals Post-PH

Next we tested if reexpression of HNF4α in hepatocytes by intravenous injection of AAV8-CMV-HNF4α could rescue HNF4α-KO mice after PH by restoring HNF4α transcriptional activity after cell division. First we reexpressed HNF4α in HNF4α-KO mice and measured HNF4α target gene expression. HNF4α reexpression increased total c-MET and total EGFR protein expression to WT levels and reduced cyclin D1 expression to WT levels (Fig. 6A). Hepatocyte proliferation was assessed by immunohistochemical analysis of PCNA positive nuclei. High levels of hepatocyte proliferation were observed in the HNF4α-KO group. HNF4α reexpression resulted in decreased hepatocyte proliferation compared to HNF4α-KO mice (Fig. 6B,C). Finally, we reexpressed HNF4α in HNF4α-KO mice 2 days post-PH. This treatment improved survival compared to HNF4α-KO mice (Fig. 6D).

**Figure 6.**
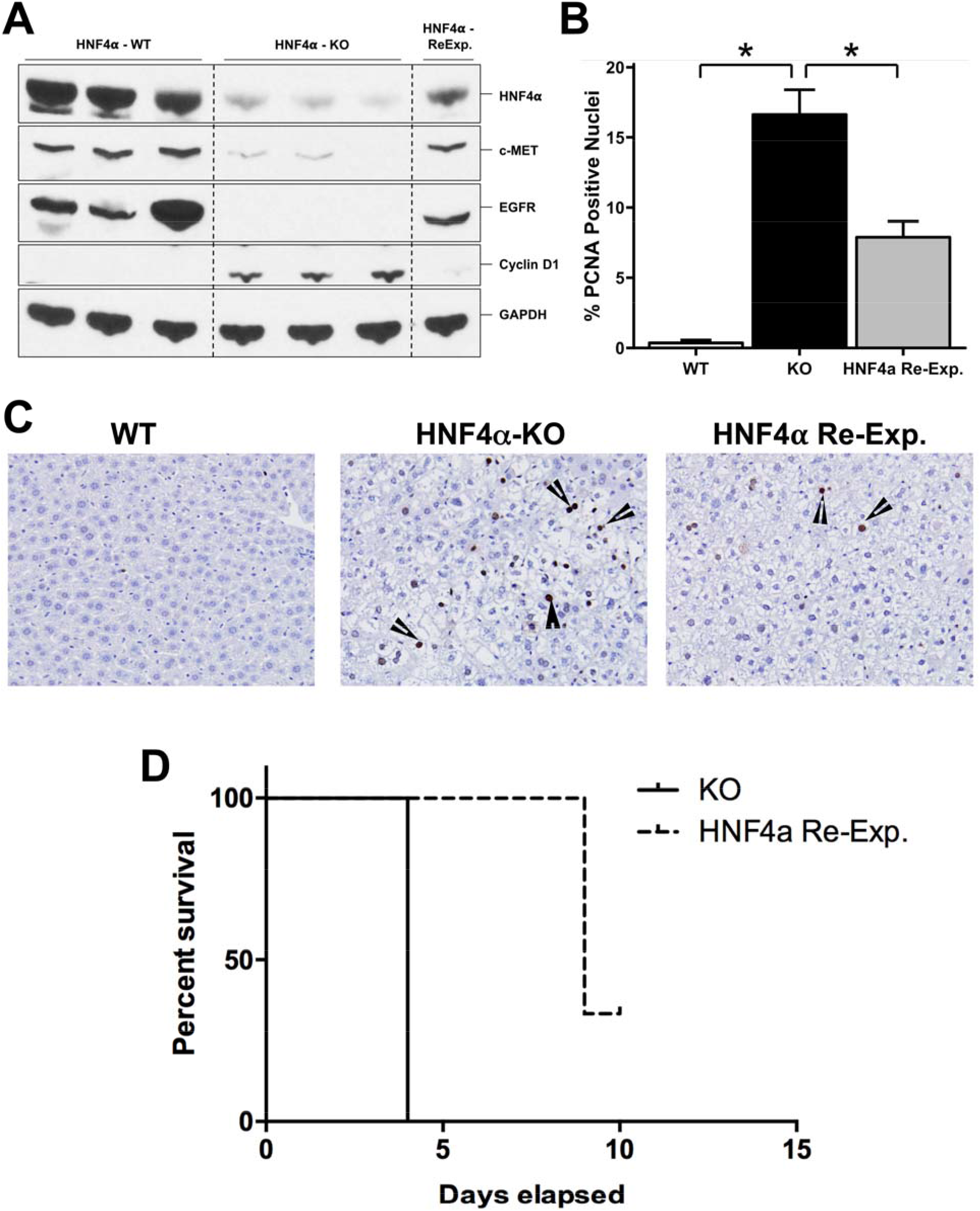
Reexpression of HNF4α restores hepatocyte quiescence and gene expression and extends survival of HNF4α-KO animals post-PH. (A) Western blot analysis of HNF4α, Cyclin D1, c-Met and EGFR in WT, HNF4α-KO and HNF4α-Reexp. mice, (B) Immunohistochemical analysis of PCNA positive nuclei in liver sections from WT, HNF4α-KO and HNF4α-Reexp. mice, (C) Representative photomicrographs (40x) of PCNA-stained liver sections, (D) Kaplan-Meier survival analysis of WT and HNF4α-Reexp mice after PH. *indicate significant difference at P ≤ 0.05.

## Discussion

HNF4α is the master regulator of hepatocyte differentiation because it is required for hepatocyte differentiation during embryonic liver development and it maintains hepatocyte specific gene expression in adult hepatocytes (3–5). HNF4α also exerts strong anti-proliferative effects on hepatocytes and decreased HNF4α expression and activity results in loss of quiescence, which can lead to HCC (14, 17, 18). Despite being recognized as one of the strongest anti-proliferative proteins in the liver, the role of HNF4α during liver regeneration is not known. We investigated the role of HNF4α in liver regeneration after PH, where adult hepatocytes exit the physiological quiescent state and enter the cell cycle before returning to the quiescent state once liver regeneration has been completed. We hypothesized that decreased HNF4α expression and function would occur during the initiation of regeneration to alleviate the antiproliferative effects of HNF4α as quiescent hepatocytes enter the cell cycle during the initiation of liver regeneration. Indeed, we observed decreased nuclear and cytoplasmic levels of HNF4α occurring within hours after surgery. This decreased protein expression was not due to changes in transcription of the HNF4α gene because there were no changes in HNF4α mRNA post-PH. Decreased periportal HNF4α staining after PH has been previously reported (38). However, HNF4α expression during liver regeneration has never been studied in a time course at this level of resolution until now. Posttranslational modifications (PTMs) of HNF4α which result in decreased nuclear localization are known (39, 40), and some PTMs can lead to proteosomal degradation of HNF4α (41). Future studies are needed to investigate whether these posttranslational modifications, especially phosphorylation of HNF4α, is the mechanism behind the decrease in HNF4α protein levels during the initiation of liver regeneration. Decrease in expression of HNF4α protein after PH resulted in decrease in its transcriptional activity as measured by its target genes, which play a critical role in liver physiology (14). Our observations independently reproduce a microarray study describing decreased HNF4α target gene expression 4 hours post-PH (42). This highlights functional differences between quiescent and proliferating hepatocytes and suggests restoring HNF4α in hepatocytes responding to chronic injury could successfully restore hepatic function (9, 43).

Next, we hypothesized that overexpressing HNF4α in hepatocytes at time points after PH when HNF4α levels were decreased would prevent hepatocytes from proliferating and halt or delay initiation of liver regeneration. Interestingly, our experiments with Tet-On driven HNF4α expression revealed no significant effect on overall hepatocyte proliferation after PH. These data suggest that rapid down regulation of HNF4α after PH may accelerate hepatocyte cell cycle entry, but is not absolutely necessary for initiation of cell cycle progression after PH. It is also possible that HNF4α overexpression resulted in delayed initiation because the mechanism responsible for HNF4α degradation was still active and more time was needed to degrade the overexpressed protein. Inhibiting the mechanism responsible for HNF4α degradation could be a better strategy to study the effects of increased HNF4α expression during initiation of liver regeneration once those mechanisms are established. One interesting observation from the HNF4α overexpression study was the lack of steatosis in HNF4α overexpressing mice, which is known to occur after PH (44). These data indicate that the initial loss of HNF4α could be mechanistically involved in the transient steatosis observed following PH and those mechanisms may be independent of initiation of cell proliferation.

The most striking observation from our studies is 100% mortality in HNF4α-KO mice following PH within 11 days secondary to loss of hepatic function. Several HNF4α-KO mice exhibited signs of hepatic encephalopathy such as lack of righting reflex prior to death (data not shown). Decreased hepatic function further manifested as elevated bilirubin levels in HNF4α-KO mice after PH. The fact that HNF4α-KO hepatocytes were not able to compensate for loss of HNF4α function (as measured by target gene expression) further support the role of HNF4α as a master regulator of hepatic differentiation. Consistent with our previous findings that HNF4α is antiproliferative in hepatocytes, the HNF4α-KO mice showed a consistently higher cell proliferation, which never declined after PH. But the lack of HNF4α resulted in failure to redifferentiate the newly divided cells resulting in significant decline in hepatic function leading to death. This was evident by the complete lack of hepatocyte differentiation genes at 7 days post PH in HNF4α-KO mice as compared to the WT mice. This was consistent with the significant increase in nuclear HNF4α expression observed in the control mice (Fig. 1) at days 3 and 5 after PH, the time points when the cell proliferation decreases and redifferentiation starts. Taken together, these data clearly demonstrate that HNF4α-mediated differentiation of the newly divided hepatocytes is an essential part of the termination of regeneration and failure to redifferentiate can lead to cell and animal mortality.

Another important observation in our studies was the complete lack of MET and EGFR expression in HNF4α-KO mice. It is known that signaling induced by primary hepatic mitogens HGF and EGF family members (EGF, TGFα etc.) via their cognate receptors c-MET and EGFR is essential for liver regeneration after PH (24). However, HNF4α-KO mice seem to be an exception to this rule. We observed complete loss of c-MET and EGFR expression in HNF4α-KO mice, yet hepatocyte proliferation occurred in these mice at high levels before and throughout liver regeneration. To address the alternate mechanisms at play, we investigated several known pathways including Wnt/β-catenin pathway and MAPK signaling. Whereas, significantly higher β-catenin signaling was observed in HNF4α-KO mice as evident by decrease in phosphorylated (inactive) β-catenin, this was observed only during the early time points. Similarly, the AKT, p38 and ERK-1/2 pathways also revealed only moderate differences between WT and HNF4α-KO mice after PH, which can not explain the sustained proliferation of hepatocytes in HNF4α-KO mice. However, we observed a significant activation of c-myc in HNF4α-KO mice consistent with the previous studies (14). Also, the role of myc in liver regeneration and cancer is well documented. Our studies show that HNF4α deleted hepatocytes lose two of the primary mitogenic pathways, namely MET and EGFR, but proliferation continues mainly via myc overexpression.

The RNAseq analysis further revealed mechanisms that drive continued proliferation and also the mechanisms that lead to liver failure and death of the HNF4α-KO mice after PH. We investigated days 2 post-PH, the time points with peak proliferation, and 5 days post-PH the time point when termination of regeneration begins. At both time points, the HNF4α-KO livers exhibited a proinflammatory and profibrotic transcriptional profile. Elevated inflammation in HNF4α-KO mice was not caused by hepatocellular injury, as there was no difference in ALT levels or histopathological changes between groups (data not shown). However, previous studies have reported elevated bile acid levels in HNF4α-KO livers, which could cause increased inflammatory signaling (45). Furthermore, many HNF4α negative target genes such as EGR1 are pro-inflammatory. IPA analysis also supported our findings of dedifferentiation and loss of hepatocyte function in HNF4α-KO mice. IPA predicted consistent inhibition of several prominent hepatocyte functions in HNF4α-KO at day 2 and day 5 post-PH. However, we observed a shift in the characteristics of activated functions between the two time points. Most of the activated functions in HNF4α-KO mice at day 2 included those involved in inflammation. However, the activated functions in HNF4α-KO mice at day 5 included many more functions involved in proliferation, tumorigenesis and a developmentally immature phenotype in addition to inflammation. Furthermore, HNF4α-KO mice at day 5 were predicted to express higher levels of known oncogenes (TRIM24, SNAI2, PLAG1) which were not activated at day 2. This transformation from an abnormal regenerative response to the beginnings of a pathological condition raise the question of how hepatocytes with poor expression of HNF4α may respond to chronic or intermittent low level liver injury. The decreased expression of HNF4α expressed in many liver diseases and HCC may be an example of the innate regenerative capability of the liver contributing to the progression of liver disease.

We observed that reexpression of HNF4α in HNF4α-KO mice using the AAV8 resulted in restoration of majority of the WT gene expression pattern. However, when HNF4α construct was introduced after PH, we did not observe complete prevention of death. This may be due to several reasons including the timing of HNF4α reexpression (two days after PH may be too early), the possibility that dividing cells may have decreased sensitivity to AAV8 and that the dividing cells may reject AAV8-mediated introduction of exogenous material. Nevertheless, we did observe a significant increase in survival following HNF4α reintroduction in HNF4α-KO mice.

In summary, our studies are the first to examine and manipulate HNF4α expression over multiple time points throughout the initiation, progression and termination of liver regeneration after partial hepatectomy. Our findings suggest downregulation of HNF4α may contribute to hepatocyte cell cycle entry during initiation of regeneration. More importantly, reestablishment of HNF4α activity during termination of regeneration is absolutely required for termination of regeneration and resumption of hepatocyte function. This study uncovers new evidence of HNF4α mediated expression of EGFR and c-MET and demonstrate that HNF4α-KO mice are capable of mounting a regenerative response despite lacking these receptors. Furthermore, regeneration in the absence of HNF4α results in a dedifferentiated, pro-carcinogenic hepatocyte phenotype. These results confirm the role of HNF4α as a major player in hepatocyte proliferation and differentiation during liver regeneration.

## Notes

**Financial Support:** These studies were supported by NIH-COBRE (P20 RR021940-03, P30 GM118247), NIEHS Toxicology Training Grant (T32ES007079-34) and NIH R01DK 0198414

## References

1. Parviz F, Matullo C, Garrison WD, Savatski L, Adamson JW, Ning G, Kaestner KH, et al Hepatocyte nuclear factor 4alpha controls the development of a hepatic epithelium and liver morphogenesis. Nat Genet 2003;34:292–296.

2. Duncan SA, Manova K, Chen WS, Hoodless P, Weinstein DC, Bachvarova RF, Darnell JE, Jr. Expression of transcription factor HNF-4 in the extraembryonic endoderm, gut, and nephrogenic tissue of the developing mouse embryo: HNF-4 is a marker for primary endoderm in the implanting blastocyst. Proc Natl Acad Sci U S A 1994;91:7598–7602.

3. Kyrmizi I, Hatzis P, Katrakili N, Tronche F, Gonzalez FJ, Talianidis I. Plasticity and expanding complexity of the hepatic transcription factor network during liver development. Genes Dev 2006;20:2293–2305.

4. Morimoto A, Kannari M, Tsuchida Y, Sasaki S, Saito C, Matsuta T, Maeda T,et alAn HNF4alpha-microRNA-194/192 signaling axis maintains hepatic cell function. J Biol Chem 2017;292:10574–10585.

5. Gonzalez FJ. Regulation of hepatocyte nuclear factor 4 alpha-mediated transcription. Drug Metab Pharmacokinet 2008;23:2–7.

6. Hu X, Xie P, Li W, Li Z, Shan H. Direct induction of hepatocyte-like cells from immortalized human bone marrow mesenchymal stem cells by overexpression of HNF4alpha. Biochem Biophys Res Commun 2016;478:791–797.

7. Wei L, Dai Y, Zhou Y, He Z, Yao J, Zhao L, Guo Q,et alOroxylin A activates PKM1/HNF4 alpha to induce hepatoma differentiation and block cancer progression. Cell Death Dis 2017;8:e2944.

8. Yin C, Lin Y, Zhang X, Chen YX, Zeng X, Yue HY, Hou JL,et alDifferentiation therapy of hepatocellular carcinoma in mice with recombinant adenovirus carrying hepatocyte nuclear factor-4alpha gene. Hepatology 2008;48:1528–1539.

9. Nishikawa T, Bell A, Brooks JM, Setoyama K, Melis M, Han B, Fukumitsu K,et alResetting the transcription factor network reverses terminal chronic hepatic failure. J Clin Invest 2015;125:1533–1544.

10. Walesky C, Gunewardena S, Terwilliger EF, Edwards G, Borude P, Apte U. Hepatocyte-specific deletion of hepatocyte nuclear factor-4alpha in adult mice results in increased hepatocyte proliferation. Am J Physiol Gastrointest Liver Physiol 2013;304:G26–37.

11. Bonzo JA, Ferry CH, Matsubara T, Kim JH, Gonzalez FJ. Suppression of hepatocyte proliferation by hepatocyte nuclear factor 4alpha in adult mice. J Biol Chem 2012;287:7345–7356.

12. Lucas B, Grigo K, Erdmann S, Lausen J, Klein-Hitpass L, Ryffel GU. HNF4alpha reduces proliferation of kidney cells and affects genes deregulated in renal cell carcinoma. Oncogene 2005;24:6418–6431.

13. Erdmann S, Senkel S, Arndt T, Lucas B, Lausen J, Klein-Hitpass L, Ryffel GU,et alTissue-specific transcription factor HNF4alpha inhibits cell proliferation and induces apoptosis in the pancreatic INS-1 beta-cell line. Biol Chem 2007;388:91–106.

14. Walesky C, Edwards G, Borude P, Gunewardena S, O’Neil M, Yoo B, Apte U. Hepatocyte nuclear factor 4 alpha deletion promotes diethylnitrosamine-induced hepatocellular carcinoma in rodents. Hepatology 2013;57:2480–2490.

15. Michalopoulos GK. Hepatostat: Liver regeneration and normal liver tissue maintenance. Hepatology 2017;65:1384–1392.

16. Tanaka T, Jiang S, Hotta H, Takano K, Iwanari H, Sumi K, Daigo K,et alDysregulated expression of P1 and P2 promoter-driven hepatocyte nuclear factor-4alpha in the pathogenesis of human cancer. J Pathol 2006;208:662–672.

17. Hatziapostolou M, Polytarchou C, Aggelidou E, Drakaki A, Poultsides GA, Jaeger SA, Ogata H,et alAn HNF4alpha-miRNA inflammatory feedback circuit regulates hepatocellular oncogenesis. Cell 2011;147:1233–1247.

18. Cai WY, Lin LY, Hao H, Zhang SM, Ma F, Hong XX, Zhang H,et alYes-associated protein/TEA domain family member and hepatocyte nuclear factor 4-alpha (HNF4alpha) repress reciprocally to regulate hepatocarcinogenesis in rats and mice. Hepatology 2017;65:1206–1221.

19. Borude P, Edwards G, Walesky C, Li F, Ma X, Kong B, Guo GL,et alHepatocyte-specific deletion of farnesoid X receptor delays but does not inhibit liver regeneration after partial hepatectomy in mice. Hepatology 2012;56:2344–2352.

20. Yanger K, Zong Y, Maggs LR, Shapira SN, Maddipati R, Aiello NM, Thung SN,et alRobust cellular reprogramming occurs spontaneously during liver regeneration. Genes Dev2013;27:719–724.

21. Wolfe A, Thomas A, Edwards G, Jaseja R, Guo GL, Apte U. Increased activation of the Wnt/beta-catenin pathway in spontaneous hepatocellular carcinoma observed in farnesoid X receptor knockout mice. J Pharmacol Exp Ther 2011;338:12–21.

22. Apte U, Singh S, Zeng G, Cieply B, Virji MA, Wu T, Monga SP. Beta-catenin activation promotes liver regeneration after acetaminophen-induced injury. Am J Pathol 2009;175:1056–1065.

23. Das AT, Tenenbaum L, Berkhout B. Tet-On Systems For Doxycycline-inducible Gene Expression. Curr Gene Ther 2016;16:156–167.

24. Paranjpe S, Bowen WC, Mars WM, Orr A, Haynes MM, DeFrances MC, Liu S,et alCombined systemic elimination of MET and epidermal growth factor receptor signaling completely abolishes liver regeneration and leads to liver decompensation. Hepatology 2016;64:1711–1724.

25. Fabregat I, Moreno-Caceres J, Sanchez A, Dooley S, Dewidar B, Giannelli G, Ten Dijke P,et alTGF-beta signalling and liver disease. FEBS J 2016;283:2219–2232.

26. Charos AE, Reed BD, Raha D, Szekely AM, Weissman SM, Snyder M. A highly integrated and complex PPARGC1A transcription factor binding network in HepG2 cells. Genome Res 2012;22:1668–1679.

27. Alpern D, Langer D, Ballester B, Le Gras S, Romier C, Mengus G, Davidson I. TAF4, a subunit of transcription factor II D, directs promoter occupancy of nuclear receptor HNF4A during post-natal hepatocyte differentiation. Elife 2014;3:e03613.

28. Jia Y, Viswakarma N, Reddy JK. Med1 subunit of the mediator complex in nuclear receptor-regulated energy metabolism, liver regeneration, and hepatocarcinogenesis. Gene Expr 2014;16:63–75.

29. Weng M-Z, Zhuang P-Y, Hei Z-Y, Lin P-Y, Chen Z-S, Liu Y-B, Quan Z-W,et alZBTB20 is involved in liver regeneration after partial hepatectomy in mouse. Hepatobiliary &Pancreatic Diseases International 2014;13:48–54.

30. Xie Z, Zhang H, Tsai W, Zhang Y, Du Y, Zhong J, Szpirer C,et alZinc finger protein ZBTB20 is a key repressor of alpha-fetoprotein gene transcription in liver. Proc Natl Acad Sci U S A 2008;105:10859–10864.

31. Santangelo L, Marchetti A, Cicchini C, Conigliaro A, Conti B, Mancone C, Bonzo JA,et alThe stable repression of mesenchymal program is required for hepatocyte identity: a novel role for hepatocyte nuclear factor 4alpha. Hepatology 2011;53:2063–2074.

32. Juma AR, Damdimopoulou PE, Grommen SV, Van de Ven WJ, De Groef B. Emerging role of PLAG1 as a regulator of growth and reproduction. J Endocrinol 2016;228:R45–56.

33. Appikonda S, Thakkar KN, Barton MC. Regulation of gene expression in human cancers by TRIM24. Drug Discov Today Technol 2016;19:57–63.

34. Palu RA, Thummel CS. Sir2 Acts through Hepatocyte Nuclear Factor 4 to maintain insulin Signaling and Metabolic Homeostasis in Drosophila. PLoS Genet 2016;12:e1005978.

35. O’Donnell KA, Keng VW, York B, Reineke EL, Seo D, Fan D, Silverstein KA,et alA Sleeping Beauty mutagenesis screen reveals a tumor suppressor role for Ncoa2/Src-2 in liver cancer. Proc Natl Acad Sci U S A 2012;109:E1377–1386.

36. Armartmuntree N, Murata M, Techasen A, Yongvanit P, Loilome W, Namwat N, Pairojkul C,et alProlonged oxidative stress down-regulates Early B cell factor 1 with inhibition of its tumor suppressive function against cholangiocarcinoma genesis. Redox Biol 2018;14:637–644.

37. Hu Y, Guo X, Wang J, Liu Y, Gao H, Fan H, Nong X,et alA novel microRNA identified in hepatocellular carcinomas is responsive to LEF1 and facilitates proliferation and epithelial-mesenchymal transition via targeting of NFIX. Oncogenesis 2018;7:22.

38. Fukuda T, Fukuchi T, Yagi S, Shiojiri N. Immunohistochemical analyses of cell cycle progression and gene expression of biliary epithelial cells during liver regeneration after partial hepatectomy of the mouse. Exp Anim 2016;65:135–146.

39. Yokoyama A, Katsura S, Ito R, Hashiba W, Sekine H, Fujiki R, Kato S. Multiple post translational modifications in hepatocyte nuclear factor 4alpha. Biochem Biophys Res Commun 2011;410:749–753.

40. Wang Z, Salih E, Burke PA. Quantitative analysis of cytokine-induced hepatocyte nuclear factor-4alpha phosphorylation by mass spectrometry. Biochemistry 2011;50:5292–5300.

41. Chellappa K, Jankova L, Schnabl JM, Pan S, Brelivet Y, Fung CL, Chan C,et alSrc tyrosine kinase phosphorylation of nuclear receptor HNF4alpha correlates with isoform-specific loss of HNF4alpha in human colon cancer. Proc Natl Acad Sci U S A 2012;109:2302–2307.

42. Jiao H, Zhu Y, Lu S, Zheng Y, Chen H. An Integrated Approach for the Identification of HNF4alpha-Centered Transcriptional Regulatory Networks During Early Liver Regeneration. Cell Physiol Biochem 2015;36:2317–2326.

43. Yue HY, Yin C, Hou JL, Zeng X, Chen YX, Zhong W, Hu PF,et alHepatocyte nuclear factor 4alpha attenuates hepatic fibrosis in rats. Gut 2010;59:236–246.

44. Shteyer E, Liao Y, Muglia LJ, Hruz PW, Rudnick DA. Disruption of hepatic adipogenesis is associated with impaired liver regeneration in mice. Hepatology 2004;40:1322–1332.

45. Hayhurst GP, Lee YH, Lambert G, Ward JM, Gonzalez FJ. Hepatocyte nuclear factor 4alpha (nuclear receptor 2A1) is essential for maintenance of hepatic gene expression and lipid homeostasis. Mol Cell Biol 2001;21:1393–1403.

